# Cryo-EM reveals structural breaks in a patient-derived amyloid fibril from systemic AL amyloidosis

**DOI:** 10.1101/2020.10.12.332569

**Authors:** Lynn Radamaker, Julian Baur, Stefanie Huhn, Christian Haupt, Ute Hegenbart, Stefan Schönland, Akanksha Bansal, Matthias Schmidt, Marcus Fändrich

## Abstract

Systemic AL amyloidosis is a debilitating and potentially fatal disease that arises from the misfolding and fibrillation of immunoglobulin light chains (LCs). The disease is patient-specific with essentially each patient possessing a unique LC sequence. In this study, we present the first ex vivo fibril structures of a λ3 LC. The fibrils were extracted from the explanted heart of a patient (FOR005) and consist of 115 residues, mainly from the LC variable domain. The fibril structures imply that a 180° rotation around the disulfide bond and a major unfolding step are necessary for fibrils to form. The two fibril structures show highly similar fibril protein folds, differing in only a 12-residue segment. Remarkably, the two structures do not represent separate fibril morphologies, as they can co-exist at different z-axial positions within the same fibril. Our data imply the presence of structural breaks at the interface of the two structural forms.

## Introduction

Systemic AL amyloidosis is a protein misfolding disease that is newly diagnosed in 4-15 persons per one million per year in the United States of America^1^ and other parts of the world^2,3^. The amyloid deposits underlying this disease frequently occur in heart and kidneys, with cardiac involvement being the most important prognostic factor for patient survival^4^. The LC amino acid sequence is extraordinarily variable, as a consequence of the recombination of different variable (V), joining (J) and constant (C) germ line (GL) segments as well as somatic hypermutation^5^. Out of the resulting LC variants, the subtypes λ1, λ2, λ3, λ6 and κ1 are in particular associated with AL amyloidosis^6,7^.

It is well established that amyloid fibrils and other LC aggregates play a defining role in the pathogenicity of this disease^4,5^. However, except for the chemotherapeutic removal of the pathogenic plasma cell clone, no pharmacological treatment options exist which directly prevent fibril formation or reverse fibril-induced organ damage^8^. One reason for this paucity of treatment options is the lack of knowledge about the mechanism of LC misfolding and the structure of pathogenic amyloid fibrils in vivo. Another reason is the patient-specific nature of systemic AL amyloidosis, with each patient presenting an essentially unique LC precursor and fibril protein^9^.

To provide insight into the fibril structure and LC misfolding mechanism in vivo, we recently set up a research strategy in which AL amyloid fibrils were extracted from diseased tissue and subjected to biochemical analysis^10,11^. The fibril proteins are mainly derived from the LC variable (V_L_) domain of the fibril protein precursor^10,11^, consistent with earlier observations^12^. They contain the intramolecular disulfide bond that is also present within the natively folded V_L_ domain^10,13^. The fibrils are polymorphic^11^, but consistent amyloid fibril morphologies are found in different organs/deposition sites within the same patient^10^. Different AL patients present different fibril morphologies^10,11^, suggesting that the variability of the LC sequence leads to different, or even patient-specific fibril structures.

To obtain insight into their molecular conformations, we and others recently started to employ cryo-electron microscopy (cryo-EM) combined with three-dimensional (3D) reconstruction. So far, two AL amyloid fibrils were analyzed with this combination of methods, one derived from a λ1 [13], termed hereafter FOR006, and one from a λ6 LC^14^. The two fibril proteins showed markedly different folds, and their conformations differed profoundly from natively folded LCs. In the present study, we analyze the structure of fibrils that were purified from the heart muscle tissue of a patient (FOR005) with λ3 LC-derived amyloid fibrils. Using cryo-EM, we obtained two different fibril structures, termed here A and B. The two structures co-exist at different z-axial positions within the same fibril, which implies the presence of structural breaks in these patient-derived amyloid fibrils.

## Results

### Extraction of the fibril protein and sequence analysis

The analyzed AL amyloid fibrils were extracted from the explanted heart of a female patient (FOR005) with systemic AL amyloidosis. The patient suffered from severe cardiomyopathy and underwent heart transplantation at the age of 50 years. We previously obtained the amino acid sequence of the fibril protein by protein sequencing, and the nucleotide sequence of the precursor LC by cDNA sequencing^10^. The tissue-deposited fibril protein corresponds to residues Ser2-Ser116 of the λ3 precursor LC and consist of the V_L_ domain and a few residues (Gly109-Ser116) of the LC constant (C_L_) domain. Bioinformatic analysis of the LC cDNA sequence indicated that it originates from the GL segments *IGLV3-19*01*, *IGLJ2*01* and *IGLC2*, which agrees with previous data showing that the *IGLV3-19* GL segment is linked to heart involvement^6,7^. The amino acid sequence of the fibril protein differs from the protein sequence of the translated GL segments in several positions, probably as a result of the B-cell clone specific somatic hypermutation. The fibril protein sequence contains five mutations with respect to the GL protein sequence within the *IGLV3-19* segment (Tyr31Ser, Tyr48Phe, Gly49Arg, Asn51Ser, Gly94Ala), one within *IGLJ2*01* (Val97Gln) and one within *IGLC2* (Val135Gly). C_L_ mutations are rarely reported for patients with AL amyloidosis, possibly because the cDNA-based gene sequencing of the fibril protein precursor is often confined to the V_L_ domain.

### Observation of two fibril structures in the fibril extracts

The extracted fibrils were subjected to cryo-EM and imaged at 300 kV (Supplementary Figure 1a). Visual inspection of the recorded images revealed, despite evidence for polymorphism^10^, one apparently dominant fibril morphology that corresponded to more than 95 % of fibrils visible in the fibril extracts. Picking this fibril morphology for 3D reconstruction and performing two dimensional (2D) and 3D classification resulted in two 3D classes showing two different fibril structures, termed here A and B (Fig. 1, Supplementary Figure 1b). The corresponding reconstructions were refined to spatial resolutions of 3.2 Å for fibril structure A and 3.4 Å for B (Supplementary Table 1), based on the 0.143 Fourier shell correlation (FCS) criterion (Supplementary Figure 2a). Their local resolution varied in the fibril cross-sections, with higher resolution occurring at the fibril center and lower resolution towards the edges (Supplementary Figure 2b). Additional rounds of 3D classification did not further subdivide the data sets in a meaningful fashion.

**Fig. 1.**
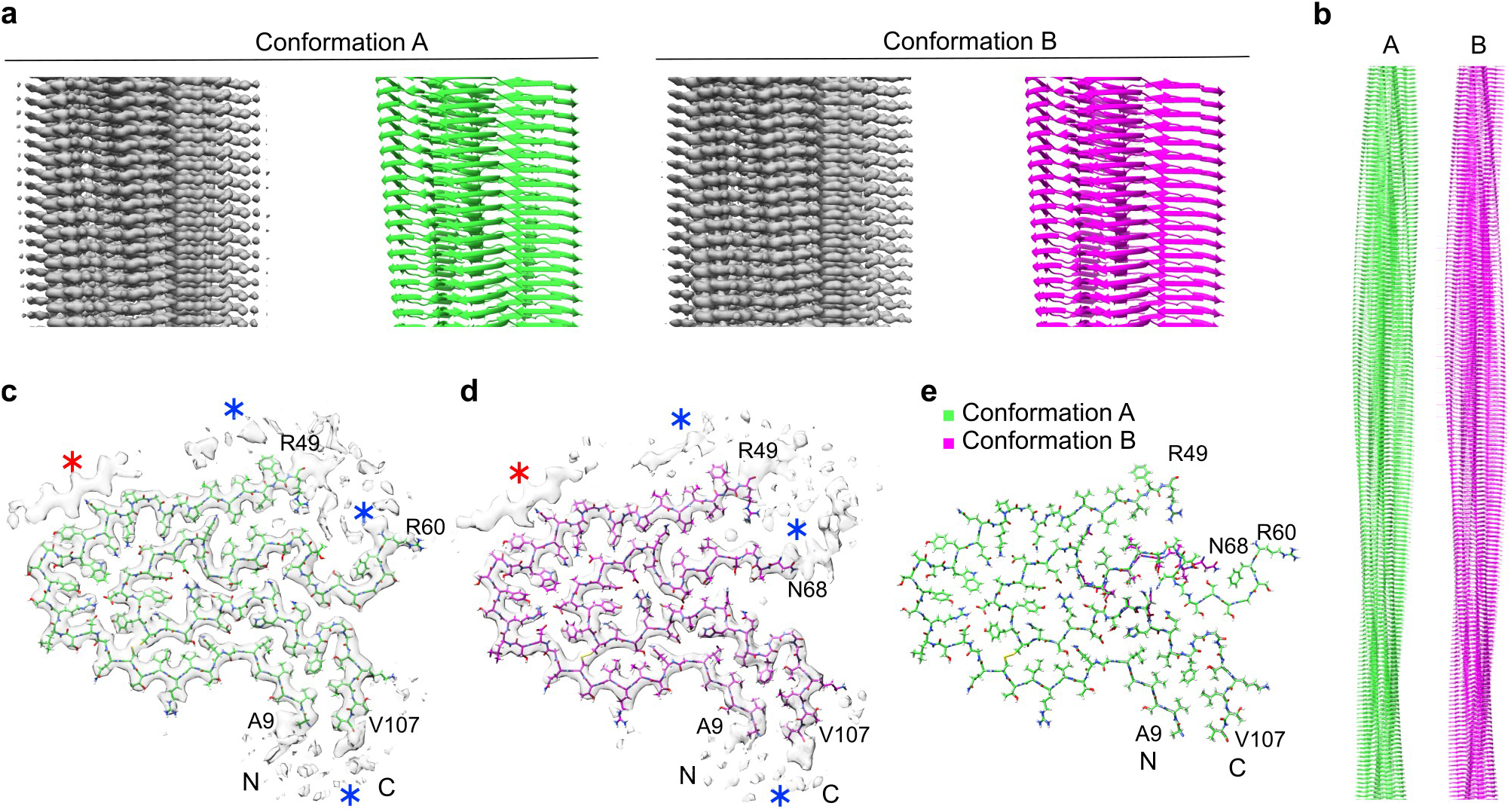
Two different fibril protein conformations underlie the FOR005 amyloid fibrils. (a) Side views of the 3D maps of fibril structures A and B (left, gray) and corresponding molecular models (right, green/magenta). (b) Side view of longer segments of the two molecular models. (c, d) Cross-sectional views of the fibril protein conformations A (c) (EMD-11031) and B (d) (EMD-11030). Blue asterisk: region with blurry density surrounding the fibril core. Red asterisk: extra density decorating the fibril core, indicating an ordered peptide conformation. (e) Overlay of the molecular models of fibril structures A (PDB: 6Z10) and B (PDB: 6Z1I). The N- and C-terminal residues of the model are highlighted.

After the initial 3D classification, the data set contained 64,652 segments classified as fibril structure A and 36,667 as fibril structure B. The final reconstructions contained 11,003 segments for fibril structure A and 12,122 for fibril structure B (Supplementary Table 1). We interpreted the two reconstructions with molecular models (Fig. 1c,d) and obtained model resolutions of 3.1 Å for reconstruction A and 3.2 Å for B (Supplementary Table 1, Supplementary Figure 2c). 2D projections of the models correspond well to the 2D class averages of the original segments (Supplementary Figure 3). Both models depict polar fibrils with C1 helical symmetry (Supplementary Table 1), consisting of only one protofilament and a single stack of fibril proteins (Fig. 1b). All peptide bonds of the fibril proteins, including the two X-Pro bonds, are modeled as *trans* isomers.

Both 3D maps contain diffuse density decorating the ordered fibril core (Fig. 1c, d, blue star), reminiscent of the two previously reported cryo-EM structures of ex vivo AL amyloid fibrils^13,14^. These diffuse density regions may represent disordered parts of the fibril protein or non-fibril components. In addition, there is a well-defined density feature (Fig 1c, d, red star) that appears to stem from a peptide segment in β-sheet conformation, owing to the zig-zag pattern and a 4.8 Å rise along the fibril axis (Supplementary Figure 4a, b). Similar well-defined density islands were previously observed with in vitro formed fibril structures^15,16^. One study suggested that the density islands were formed from a segment of the fibril protein that was protruding from the main fibril core^15^. In another study the density island originated from a peripherally attached fibril protein that adopted a single, short cross-β strand but that was otherwise conformationally disordered^16^. As all segments outside the FOR005 fibril core are too short to reach our density islands (Supplementary Figure 4c), we conclude that non-covalently attached fibril proteins, or fibril protein fragments, are the most plausible explanation of the density islands in the FOR005 fibril structure.

### Comparison of the fibril protein conformations A and B

The two fibril structures arise from similar but slightly different protein conformations. The fibril proteins are essentially indistinguishable at residues Ala9-Arg49 and Leu72-Val107. Residues Ser2-Pro8 and Leu108-Ser116 could not be assigned to any well-defined density in the 3D map, which implies that these segments are structurally heterogeneous or disordered. The main difference between the two structures lies in the segment Arg60-Ser71 (Fig. 1e). In fibril A, residues Arg60-Ser71 are in a stable conformation encompassing an arch, while residues Lys50-Asp59 are not well defined in the 3D map (Fig. 1c). In fibril B, residues Asn68-Ser71 are in a relatively extended conformation and the region of structural disorder occurs between residues Lys50-Gly67 (Fig. 1d). Importantly, mass spectrometry previously demonstrated the fibril protein to be continuous and to extend from Ser2 to Ser116 [10]. Thus, the fibril core as seen in our 3D map is not made up of two fibril protein fragments, but instead it consists of two structurally ordered segments (Ala9-Arg49 and Arg60/Asn68-Val107) that are linked by a structurally heterogeneous region (Lys50-Asp59/Gly67).

The fibril protein shows β-strand conformation at residues Val10-Leu14, Thr17-Gln23, Asp25-Ser26, Arg28-Ser31, Trp34-Gln37, Pro43-Ile47, Leu72-Thr75, Ala79-Glu82, Tyr85-Tyr86, Asn88-Asp91, Asn95-Gln97, and Thr103-Thr106 in both fibrils (Fig. 2a). We refer to these segments as β1 to β12. Structure A contains two additional β-strands in a segment that is disordered in structure B (Arg60-Gly67). These strands are formed by residues Arg60-Ser62 and Ser64-Ser65 and are termed β6’ and β6’’ because they are in between the strands β6 and β7. All strands form cross-β sheets with parallel, hydrogen bonded strand-strand interactions (Fig. 2b, Supplementary Figure 5a). The protein fold is compact and devoid of large internal cavities.

**Fig. 2.**
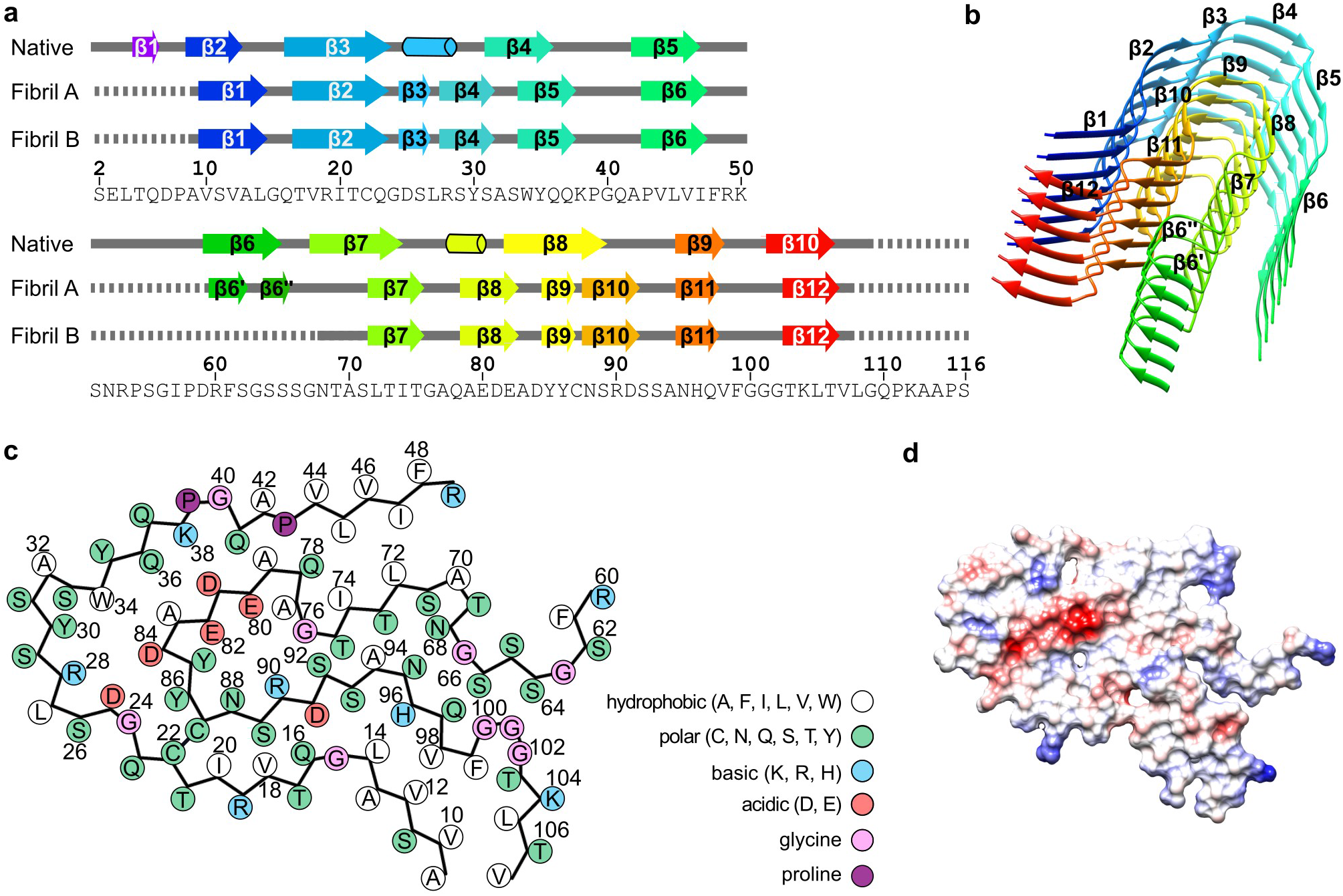
Secondary structure and folding of the fibril proteins. (a) Schematic representation of the secondary structures of the fibril proteins A and B, and of a crystal structure of the refolded fibril protein (native, PDB: 5L6Q10). Arrows indicate β-strands and cylinders α-helical conformations. Continuous lines indicate ordered conformation, dotted lines indicate disordered segments. The definition of secondary structural elements follows the definition in the respective manuscripts. (b) Ribbon diagram of a stack of six fibril proteins (conformation A). (c) Schematic representation of the amino acid positions in conformation A. (d) Electrostatic surface representation of the fibril protein conformation A. Red indicates negative charge, blue positive and white neutral. Supplementary Figure 5 shows the corresponding images for fibril conformation B.

### Molecular interactions defining the fibril structure

The fibril proteins interact along the fibril axis through backbone hydrogen bonds that extend between the strands of the cross-β sheets. In addition, there are side chain-side chain interactions, such as polar ladders of asparagine or glutamine residues or stacked hydrophobic or aromatic groups. These features are shown for residues Gln37 and Phe48 in the Supplementary Figure 6a, b. The fibrils show axial height changes of 7.8 Å (fibril A) and 7.1 Å (fibril B), which lead to polar fibril topologies and sterically interlock the fibril layers. Each fibril protein layer interacts only with the layers above and below, e.g. Lys38 from layer *i* interacts with Asp81 from layer *i*+1 (Supplementary Figure 6c).

The surfaces of both fibrils are rich in charged and polar amino acids (Fig. 2c, Supplementary Figure 5b). The fibril cores contain small hydrophobic patches, such as the one formed by residues Val10, Val12, Leu14, Val98, Phe99, Leu105 and Val107 (Fig. 2c, Supplementary Figure 5b), as well as patches of buried polar residues. The structure buries a number of compensating charge-charge interactions, for example at residues Asp25 and Arg28, Arg28 and Asp84, Lys38 and Asp81, Glu80 and Arg90 (Fig. 2c, Supplementary Figure 5b), as well as an acidic moiety, which is not fully charge compensated. This moiety is formed by residues Glu80, Asp81, Glu82 and Asp84 (Fig. 2d), resembling the partially uncompensated acidic moiety in the previously described λ1 fibril structure^13^. In contrast to the previous λ1 fibril, however, there is no water-filled cavity around the acidic moiety in our fibril.

### Location of aggregation-prone segments and mutations

The mutagenic changes of the amyloidogenic LCs compared with the GL sequences are widely believed to trigger amyloidosis in the respective patients^9,17^. However, analysis of the mutated positions within our structure does not readily offer an explanation for their pathogenicity. Some mutations, such as Asn51Ser and Val135Gly, lie within a part of the precursor protein that is disordered or cleaved off in the fibril (Fig. 2a). In addition, none of the mutations affecting the fibril core adds an obviously favorable interaction. The non-conservative Gly49Arg mutation even leads to a buried charge that is not compensated by a nearby opposite charge (Fig. 3a), suggesting that this mutation may even be unfavorable to the fibril structure. Moreover, analysis of the location of the mutations within the native LC does not readily provide evidence that they might be destabilizing to the native protein conformation (Supplementary Figure 7a, b). The mutations do not remove an obviously stabilizing interaction and do not affect internal residues that might be considered to be crucial for protein stability. Instead, all mutations are located on the surface of the globularly folded LC (Supplementary Figure 7a, b).

**Fig. 3.**
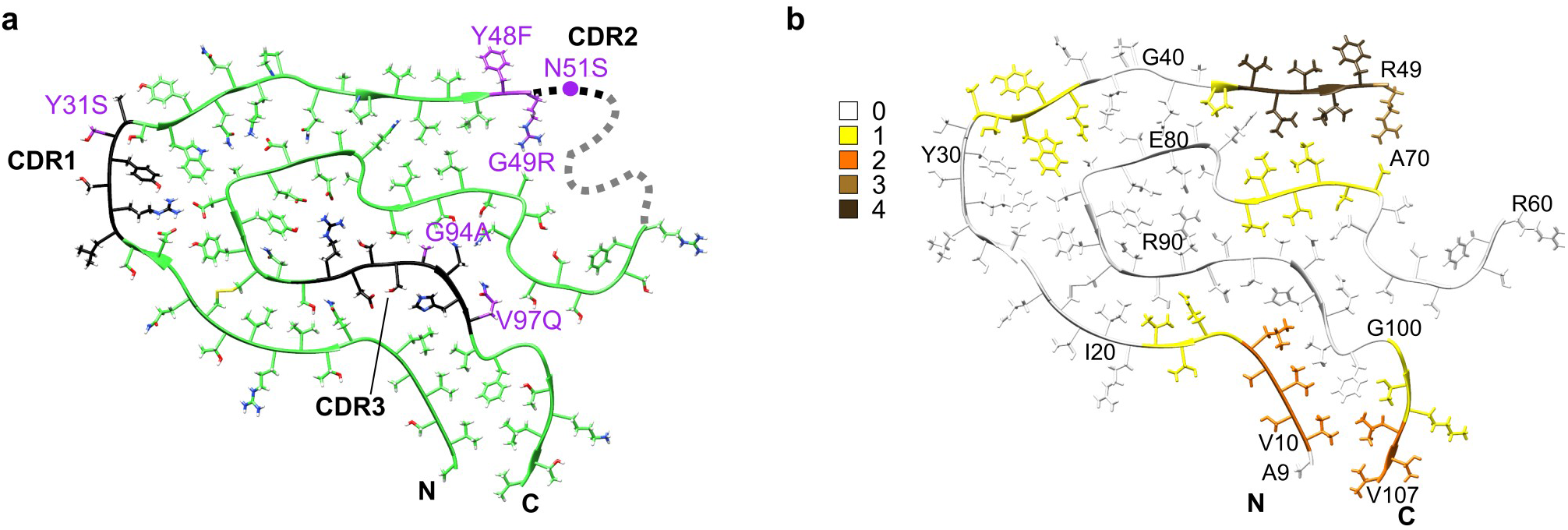
Location of the mutational positions and aggregation prone regions in the fibril structure. (a) Mutations with respect to the GL protein sequence (purple) and CDRs (black) marked in fibril structure A. The dotted line represents the disordered segment. (b) Molecular model of conformation A colored according to aggregation score.

Computer-based predictions of the aggregation propensity of the FOR005 LC identified the highest aggregation score in the V_L_ domain at residues Val44–Arg49 (Fig 3b, Supplementary Figure 8). These residues form a hydrophobic patch on the fibril surface, which is decorated with the extra density region described above (Fig. 1c, d, blue star). The three disordered protein segments in the fibril protein (Ser2-Pro8, Lys50-Gly67, Leu108-Ser116) correlate with regions with low aggregation scores (Supplementary Figure 8). However, comparing the aggregation score of the FOR005 LC to that of the GL protein sequence (Supplementary Figure 8) does not reveal any clear trend whether the FOR005 LC or its putative GL precursor is more aggregation prone (Supplementary Figure 8). We conclude that the effect of mutation is subtle and not readily evident by the above analysis. Support for this view comes from a recent study in which the rather counterintuitive observation was reported that a conservative leucine to valine mutation on the surface of a patient-derived V_L_ domain is strongly destabilizing to the native protein structure and promotes the formation of amyloid fibrils in vitro^18^.

### Conformations A and B co-exist within the same fibrils

Finally, we sought to determine whether the two reconstructed 3D maps A and B represent two different fibril morphologies or whether the two structures co-exist within the same fibril particle. By visual inspection of the cryo-EM micrographs and measurement of global parameters, such as fibril width or cross-over distance, we could not categorize the fibrils in our sample into separate structures A and B. Also, the reconstructed 3D maps have identical helical parameters, such as fibril symmetry, polarity, axial rise and twist value, as well as a fibril pitch of 155 nm (Supplementary Table 1). The difference between the two structures could only be revealed when the fibril images were cropped into segments that were then aligned independently of their structural context during 3D classification.

Analyzing the origin of the fibril segments in the respective data sets producing reconstructions A and B we would have expected, for separate morphologies, that each fibril contains segments belonging to only one of the two data sets A or B (except for minor classification errors). Surprisingly, however, we found that most fibrils in our sample showed a mixture of A and B segments (Fig. 4a, Supplementary Figure 9a). Fractions of type B segments per fibril varied almost continuously from 0 to 1 (Fig. 4b, Supplementary Figure 9b). The assignment of segments to fibril protein conformations A and B could not readily be correlated with certain positions on the fibril helix on the cryo-EM micrographs, for example the cross-over or the segment in between two cross-overs, which might have suggested problems in their alignment. Moreover, the segments are not randomly distributed across the fibrils but mostly separated into distinct regions along the fibril axis in which all segments correspond to either structure A or B (Fig. 4a). These results were obtained consistently across different data sets, including the data set resulting from the initial 3D classification (Fig. 4a, b) as well as the data set from the final reconstructions (Supplementary Figure 9a, b). In conclusion, the fibrils in our data set cannot simply be divided into two fibril morphologies A and B. Instead, the two fibril structures A and B occur simultaneously within a fibril protein stack. This observation indicates that there are structural breaks at the interface of fibril regions corresponding to structures A or B (Fig. 4c, d).

**Fig. 4.**
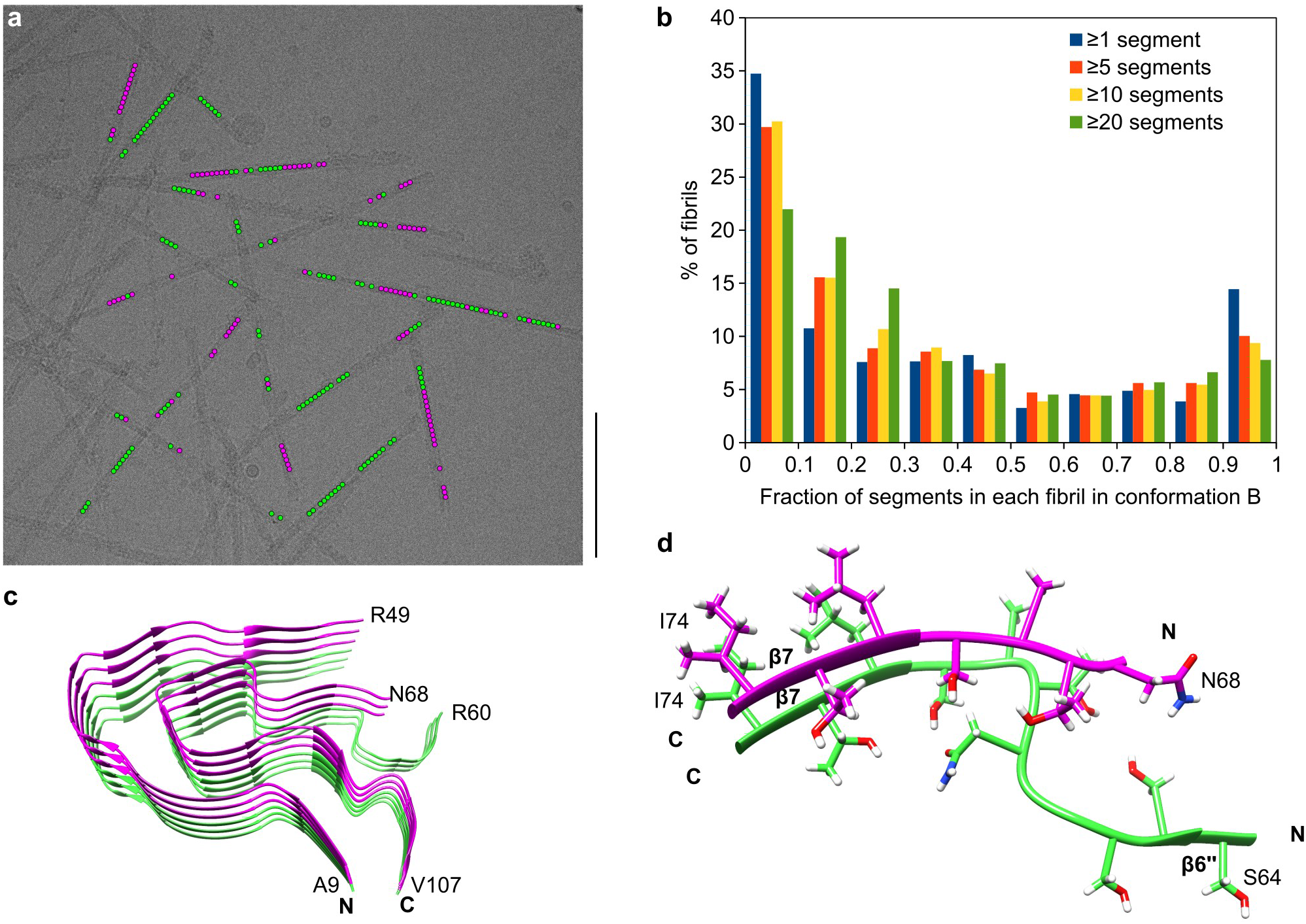
Evidence for structural breaks in FOR005 fibrils. (a) Representative cryo-EM micrograph showing the location of segments classified after the first 3D classification as fibril structure A (green) and fibril structure B (magenta). Scale bar: 100 nm. Supplementary Figure 9a shows the same image, highlighting only the segments used in the final fibril reconstruction. (b) Histogram of the fraction of segments classified after the first 3D classification as fibril structure B, per fibril. Four different thresholds were chosen for the minimum total number of segments per fibril: a minimum of 1 segment (11,194 fibrils), 5 segments (7,738 fibrils), 10 segments (4,278 fibrils) and 20 segments (951 fibrils). In total, the data set contained 64,652 segments classified as fibril structure A and 36,667 as fibril structure B. Supplementary Figure 9b shows an analogous histogram, but including only the segments used for the final reconstructions. (c) Stack of three fibril proteins in conformation B (magenta) on top of three fibril proteins in conformation A (green), illustrating the presence of structural breaks within the patient amyloid fibrils. (d) Detailed view of a structural break, including side chains.

## Discussion

We here present the cryo-EM structures of two amyloid fibrils (A and B) that were extracted from the explanted heart of a patient (FOR005) with systemic AL amyloidosis. The spatial resolutions are 3.2 Å for fibril structure A and 3.4 Å for fibril structure B (Supplementary Table 1). These resolutions are sufficient to establish the overall fibril topology and the fibril protein fold. However, uncertainty remains in the exact conformation of the backbone and side chains. This problem is further exacerbated by the known artefacts of cryo-EM structures, such as a loss of side chain density due to beam damage^19^, which could be relevant in our reconstructions e.g. at residues Glu80 or Lys104 (Fig 1c, d).

Systemic AL amyloidosis is a patient-specific disease^9^. Identification of common structural features in different patient-derived amyloid fibrils is potentially informative about common steps in the misfolding pathways across patient cases. Based on the available cryo-EM structures of ex vivo fibrils from systemic AL amyloidosis (Fig. 5, Supplementary Figure 10), the following commonalities can now be identified: The extracted fibril samples contain a dominant fibril morphology that consists of a single, polar protofilament. The fibril core is formed by the V_L_ domain of the precursor λ-LC in all cases. The C_L_ domain is structurally disordered and/or lost by proteolysis. The fibril proteins retain the intramolecular disulfide bond of the native V_L_ domain, indicating that LC misfolding happens in an oxidative environment, such as the extracellular space or an endocytic compartment, that retains the disulfide bond of the native V_L_ domain. The fibril proteins show an antiparallel N-to-C orientation at the disulfide that is flipped by 180° relative to the native state. The fibril protein conformations differ substantially from the natively folded LC, demonstrating that a global structural rearrangement and/or unfolding reaction takes place during the conversion of the native LC, or of a LC fragment, into a fibril. The fibrils are decorated by blurry density of uncertain origin and may arise from fibril protein segments outside the ordered core, or cellular factors attached to the fibril surface.

**Fig. 5.**
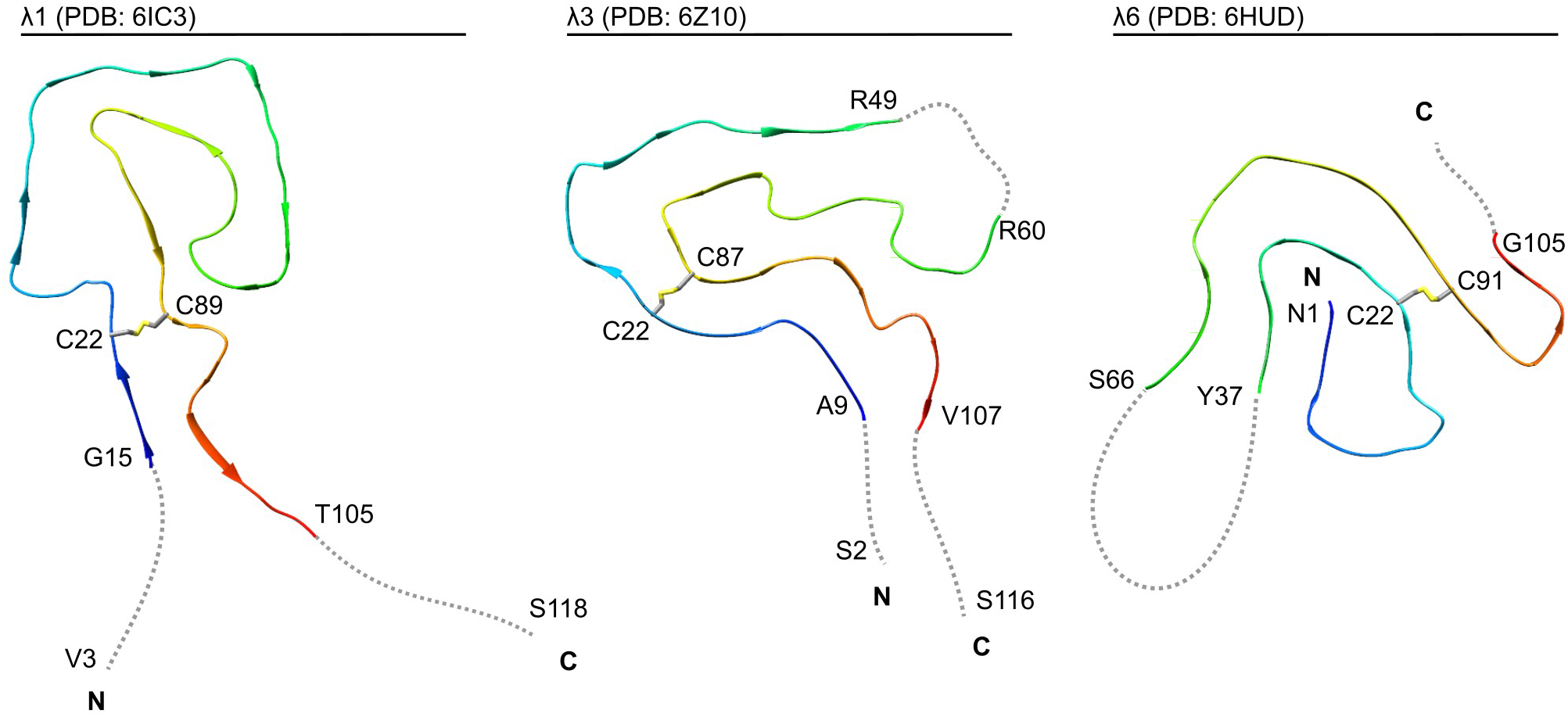
Comparison of available cryo-EM structures of ex vivo AL amyloid fibrils. Ribbon diagrams of a λ1 fibril (PDB: 6IC3^13^), the current λ3 fibril (conformation A, PDB: 6Z10) and a λ6 fibril (PDB: 6HUD^14^). The fibrils are shown in a cross-sectional view. For all structures, the location of the disulfide-bond forming cysteine residues is marked. Disordered segments are represented as dotted gray lines and depicted in arbitrary conformation. The first and the last residue of the ordered segments as well as the first and the last residue of the fibril protein are indicated, if known.

Patient-specific features of the fibrils include the exact fold of the fibril protein (Fig. 5) and the location of the β-strands and disordered segments within the sequence (Supplementary Figure 10). The current λ3- and the previous λ1-derived fibril proteins possess solvent-exposed and conformationally disordered N-termini, while the N-terminal segment of the λ6-derived fibril is buried in the fibril core and part of a β-strand (Fig. 5, Supplementary Figure 10). The C-termini are disordered in each of these fibril proteins. Our current fibril structures and the previous λ6 fibril structure each contain an internal, disordered segment interrupting the fibril protein fold. In contrast, the fold of the λ1 fibril protein is continuous (Fig. 5, Supplementary Figure 10). The λ1 fibril possesses three large channels, two of which are thought to be water-filled, while the third one contains an apolar molecular inclusion^13^. No such channels or inclusions were identified in the other fibril structures. A feature unique to the current λ3 fibrils is a well-resolved density island attached to the fibril core (Supplementary Figure 4a, b).

Our structures also differ from a number of studies which used nuclear magnetic resonance (NMR) spectroscopy to investigate the structure of LC-derived fibrils formed in vitro. These fibrils were formed from V_L_ domain constructs and include murine κ [20], human κ1 [21] as well as human λ3 [22] and λ6 sequences^23^. Particularly relevant in this case is the comparison of our structures to the NMR analysis of recombinant FOR005 V_L_ domain fibrils^22^. These fibrils were seeded in vitro with amyloid fibrils that were extracted from the heart of the patient (FOR005) with the aim to propagate the ex vivo fibril structure in the in vitro seeded fibrils^22^. Comparison of the in vitro seeded fibrils with our cryo-EM structures of patient fibrils revealed several differences. First, the in vitro seeded fibrils contained β-strand structure at residues Ile57-Pro58, which are conformationally disordered in the ex vivo fibrils (Fig. 2a). Second, the ex vivo fibrils possess a stable β-strand at residues Thr103-Thr106 that are outside the ordered core of the in vitro seeded fibrils. Third, the in vitro seeded fibrils contain a salt bridge between residues Arg49 and Asp25 [22], which are far apart in the protein fold of the patient fibril (Fig. 2c).

These data demonstrate that the in vitro fibrils are structurally different from the ex vivo fibrils analyzed here with cryo-EM. In vitro seeding with ex vivo FOR005 fibrils did not propagate, in this case, the seed structure to the daughter fibrils, although it modified the fibril structure compared with unseeded fibrils^22^. These observations imply that the seeding mechanism did not involve a replication of the seed structure. Early work with sickle cell hemoglobin identified two possible seeding mechanisms: homogeneous and heterogeneous nucleation. Homogeneous nucleation involves the attachment of the soluble fibril precursor proteins to the fibril tip, while heterogeneous nucleation involves the nucleation of new fibrils on the lateral cylindrical surface of an existing fibril^24^. While attachment of the fibril precursor protein to the tip of an amyloid fibril would be expected to lead to a replication of the fibril protein fold, this replication of the seed protein structure may not necessarily occur during heterogeneous seeding. Indeed, the observation of density islands on the outside of the FOR005 fibril core structure (Fig. 1c,d red star, Supplementary Figure 4a, b) implies the attachment of fibril proteins that do not fully replicate the fold of the fibril protein. Hence, it is important to use patient-derived fibrils when investigating the structural basis of disease. A similar conclusion was obtained previously when FOR005 fibril protein was extracted from the patient’s heart, denatured in guanidine, refolded and fibrillated in vitro (without seeds). These in vitro fibrils also showed a different morphology than the fibrils that were purified from FOR005 patient tissue^10^, as judged by transmission electron microscopy.

A particularly interesting finding in the present study is the observation of structural breaks. So far, it has been part of our general understanding of amyloid fibril structures that these are conformationally uniform along the fibril axis. Occasionally fibrils were reported that differed morphologically at its two ends^25,26,27,28^. Some of these cases could be attributed to a fibril cross-seeding, that is, the attachment of a different fibril precursor protein to a fibril tip, or to a splintering of a multi-protofilament fibril into fibril morphologies with a smaller number, or a different arrangement of protofilaments. In other cases it was unclear whether two fibril morphologies may have annealed after their formation. In our samples, however, the majority of fibrils show a mixture of conformations, and show multiple, seemingly arbitrary switching between conformations A and B in each fibril. Therefore, short segments, possibly down to a single protein layer, may be able to adopt a conformation different from that of the surrounding layers. The fibril breaks and the two fibril structures defining the breaks emerged at the 3D classification stage in our analysis and resolved a previously blurry density region into two distinct density paths (Supplementary Figure 1b). Unresolved density regions resulting from one or more disordered segments of the protein chain are reported for the majority of cryo-EM structures of in vitro and ex vivo amyloid fibrils^29^. Therefore, structural heterogeneity, such as described here, could be relevant to other fibril structures as well. Furthermore, it is possible that our fibrils contain fibril protein structures, which we were unable to resolve so far. We originally extracted 194,502 fibril segments from the cryo electron micrographs and used only 11,003 (A) and 12,122 (B) of these segments for the final 3D reconstructions (Supplementary Table 1).

Two possible scenarios can be envisioned to explain the mechanisms of the formation of structural breaks. One scenario is that they appear during fibril assembly due to an imperfect replication of the seed structure as a new molecule attaches to the fibril end (Fig. 6). Consistent with this idea, real-time microscopy studies explained the stop-and-go kinetics during fibril growth with irregularities in the addition of molecules to the tip of a growing fibril^30^. The other scenario is that breaks emerge after fibril assembly, for example because initially disordered segments adopt different stable conformations, which then proliferate along the fibril axis (Fig. 6). Support for the latter mechanism is provided by the fact that type A and B fibril proteins are mostly identical and that the differences are confined to a small segment that lies in the vicinity of an unstructured region.

**Fig. 6.**
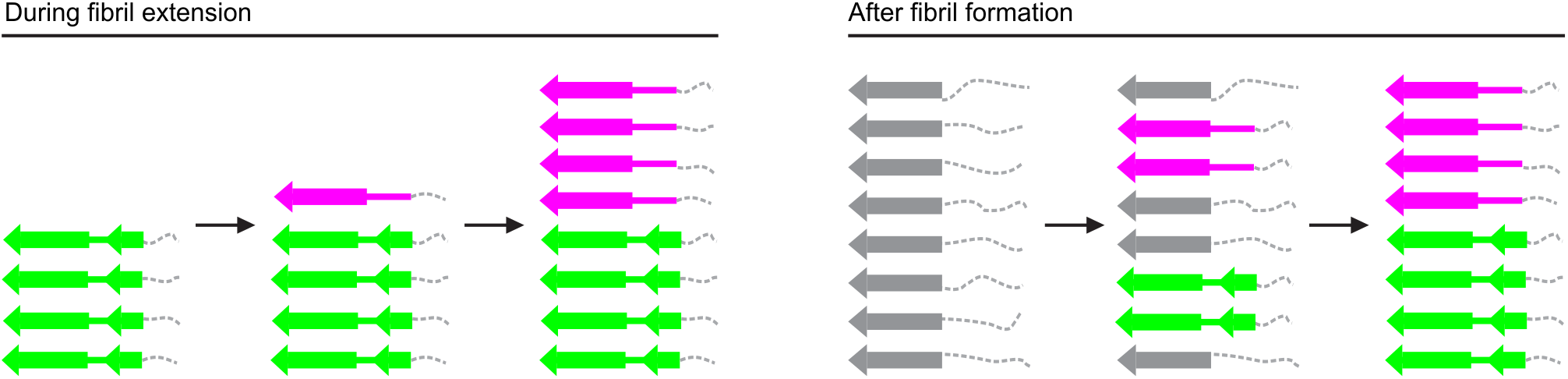
The origin of structural breaks: two possible scenarios. Schematic representation of a stack of fibril proteins, illustrating two different hypotheses on how structural breaks form: during fibril extension (left) or after fibril formation (right). Conformation A is represented by two β-sheets (green). Conformation B is represented by one β-sheet (magenta). Disordered segments are represented by dotted gray lines. Gray arrows represent the immature fibril proteins, before the mature conformations A and B are fully adopted.

While more work is necessary to discriminate between these two mechanisms, our observations lead to an important change in our understanding of the assembly of polypeptide chains into amyloid fibrils. They demonstrate that these linear aggregates are not as perfectly regular and uniform as has generally been assumed by most previous studies. Considering that the breaks were revealed in the FOR005 fibril samples only at an advanced stage of the analysis, we would predict that they will be observed more frequently in the future as the methods of structural biology become more powerful and will be able to resolve such fine details more routinely. Structural breaks and other ‘structural defects’ in cross-β sheets could have significant ramifications for the biological properties of amyloid fibrils. Examples hereof include the fragility^31^ and the loss of torsional coherence of amyloid fibrils^32,33^, the branching of amyloid fibrils during fibril outgrowth^34^ and the ability of molecular chaperones to bind to, to sever and to break down amyloid aggregates^35^.

## Methods

### Source of AL fibrils

Heart tissue was collected from a female patient (FOR005) at the age of 50, suffering from AL amyloidosis and consequent advanced heart failure. A monoclonal gammopathy was the underlying condition. The patient was treated within the heart transplant program of the University Hospital Heidelberg. The explanted heart tissue was stored at −80 °C. The study was approved by the ethical committees of the University of Heidelberg (123/2006) and of Ulm University (203/18). Informed consent was obtained from the patient for the analysis of the amyloid deposits.

### Fibril extraction from patient tissue

Applying a previously established protocol^11^ for fibril extraction, 250 mg of patient heart tissue were diced finely and 0.5 mL of ice-cold Tris Calcium Buffer (20 mM Tris, 138 mM NaCl, 2 mM CaCl_2_, 0.1 (w/v) % NaN_3_, pH 8.0) added. The sample was homogenized using a Kontes Pellet Pestle, after which it was centrifuged for 5 min at 3,100 × *g* at 4 °C. The washing step was repeated 5 times and each supernatant was stored for further analysis. Afterwards, 1 mL of freshly prepared 5 mg mL^−1^ *Clostridium histolyticum* collagenase (Sigma) in Tris calcium buffer with ethylenediaminetetraacetic acid (EDTA)-free protease inhibitor (Roche) were added and the pellet resuspended. Overnight incubation at 37 °C was followed by a 30 min centrifuge cycle at 3,100 g. Ten further washing steps with 20 mM Tris, 140 mM NaCl, 10 mM EDTA, 0.1 % (w/v) NaN_3_ and ten subsequent steps with ice cold water were then performed on the pellet, using a pipette for homogenization. One of the water supernatants was selected for cryo-EM.

### Cryo-EM

Holey carbon-coated grids (C-flat 1.2/1.3 400 mesh) were glow-discharged using 40 mA for 40 s. Using a Vitrobot (Thermo Fisher Scientific), 3.5 μL of the extracted fibril sample were incubated onL of the extracted fibril sample were incubated on each grid for 30 s at a humidity of >95 %, the excess fluid was blotted off and the grid plunged into liquid ethane, then transferred to a grid box. After plunging, one grid from each grid box (containing 4 grids) was checked using a 200 kV Jeol JEM 2100F electron microscope (Ulm University). The remaining grids in the grid boxes were kept at liquid nitrogen temperature. Cryo-electron microscopic image acquisition of one selected grid was performed using a Titan Krios transmission electron microscope (Thermo Fisher Scientific) at 300 kV equipped with a K2-Summit detector (Gatan) in counting mode. A Gatan imaging filter with a 20 eV slit was applied. The data acquisition parameters can be found in Supplementary Table 1. Global parameters of the fibril morphologies were measured using Fiji^36^. No clearly identifiable second morphology was found and the occurrence of the main morphology was estimated at over 95 %.

### Helical reconstruction

Helical reconstruction was performed using Relion 2.1 [37]. The raw data were converted from TIFF to mrcs format using IMOD^38^. Motion- and gain-correction as well as dose-weighting were performed using Motioncor2 [39]. The non-dose-weighted micrographs were used to determine the defocus values with Gctf^40^, whereas the dose-weighted micrographs were used for all subsequent steps. Fibrils were manually selected from 1,964 micrographs and segments were extracted with a box size of 300 pixels (312 Å) with an inter-box distance of 33.6 Å (~11 %). Two rounds of reference-free 2D classification were performed with a regularization value of T = 2. Class averages which showed the helical repeat along the fibril axis were selected, whereas classes showing artefacts and noise were discarded resulting in a selection of 101,319 particles. The selection was confirmed by manually arranging the class averages into a full 2D fibril side view. An initial model from a previous reconstruction was filtered to 60 Å resulting in a rod-like structure. This initial model was used as a reference to create a first 3D map using 3D classification with 553 particles picked from 9 micrographs. The resulting map was used as a reference for 3D classification with 6 classes and T = 3 using the 101,319 particles obtained from the 2D classification selection. Three of the six classes showed a clear backbone and revealed the presence of two different conformations. These classes were selected (60,044 particles) and further rounds of 3D classification with 4 classes and step-wise increasing T-values from 3 to 20 were performed. Approximately 11,003 particles were selected for conformation A and 12,122 particles for conformation B. A final round of 3D classification with increasing T-values from 80 to 200, followed by 3D auto-refinement and post-processing with a soft-edged mask and an estimated sharpening B-factor of −33.1013 Å^2^ (conformation A) and −55.1958 Å^2^ (conformation B) for each of the two conformations led to the post-processed maps. Helical z-percentages used for 3D-classification and 3D-refinement varied between 0.1 and 0.3. The twist and rise were 1.11 and 4.79 respectively, for both conformations. These values agree with the measured cross-over distances on the motion-corrected cryo-EM micrographs, and the corresponding power spectra. A left-handed twist was assumed. The resolution of each map was estimated from the value of the FSC curve for two independently refined half-maps at 0.143.

### Model building and refinement

The model was manually built using the software Coot^41^. The process was initiated by tracing a poly-Ala chain along the 3D map. The alanine residues were mutated to the LC sequence as determined previously^11^. The model obtained was subjected to manual as well as automated refinement using phenix.real_space_refine^42,43^ tool as implemented in phenix with non crystallographic symmetry and secondary structure restraints imposed. This process was repeated until a satisfactory map to model fit was obtained. This model was subsequently used to perform model based automated sharpening of the map (phenix.auto_sharpen^42,43^). The sharpened map was used for improving the model further. The final refined model was evaluated for its quality using the MolProbity^44^ generated validation report.

### Sequence analysis

The amino acid sequence of the LC investigated in this study was taken from the gene bank entry KX290463 [10], which was obtained by cDNA sequencing of FOR005 patient material. The residue numbering throughout this article refers to the precursor LC sequence (GenBank ANN81988.1) from the patient, starting with the first residue (Ser1) after the signal sequence, which is cleaved off in the fibril protein. All mutations in this manuscript are represented in the direction GL to FOR005 fibril protein. The sequence elements were defined as follows. First, the patient cDNA was translated to the putative amino acid sequence of the fibril precursor protein. Then, the cDNA of the patient and the corresponding amino acid sequence were analyzed to determine the most probable GL segments using the vbase2 [45] and BLAST/BLAT search tools [46], (http://www.ensembl.org). This analysis yielded several hits for possible GL segments. The cDNA sequences and corresponding amino acid sequences of these V, J and C GL segments were retrieved from the vbase2 [45], ENSEMBL^47^ and http://www.imgt.org^48^ databases, and genetic distances to the patient sequence were calculated by maximum composite likelihood to confirm the most probable V, J and C GL segments. Finally, the cDNA sequence of the patient and the corresponding amino acid sequence were aligned to these GL segments (fit >80%) using the MEGA multiple sequencing alignment tool “MUSCLE”^49^ and Clustal Omega (https://www.ebi.ac.uk/Tools/msa/clustalo/)^50^ to identify patient-specific mutations. The definitions of the CDRs were taken from a previous manuscript^10^.

LC aggregation propensities for every residue were calculated using the programs TANGO^51^, Foldamyloid^52^, Aggrescan^53^ and PASTA 2.0 [54]. For every prediction tool standard parameters were chosen. For Tango (Version 2.1), every amino acid which scores above 5 % was counted as a hit. For Foldamyloid, the “triple hybrid” scale was used, meaning that 5 successive amino acids with a score above 21.4 were counted as hits. PASTA 2.0 calculated aggregation with 90 % sensitivity and predicted hits if the energy cut-off fell below −2.8 PASTA Energy Units. For Aggrescan, all amino acids with values above −0.02 were counted as hot spots. In this study, an aggregation score of 0 means that none of these programs identified the corresponding residue as aggregation prone. An aggregation score of 4 means that all four programs identified the corresponding residue as prone to aggregation.

### Protein structure representation

UCSF Chimera^55^ was used for creating the images of the density maps and protein models. The structure of the refolded FOR005 fibril protein was obtained previously using protein X-ray crystallography and has the protein data bank (PDB) entry 5L6Q^10^. The native C_L_ domain is the *IGLC2* segment of PDB 4EOW^56^, where the side chain of residue Val135 was not shown in order to represent the mutated residue Gly135 of the fibril protein.

## Data availability

The reconstructed cryo-EM maps were deposited in the Electron Microscopy Data Bank with accession codes EMD-11031 (structure A) and EMD-11030 (structure B). The coordinates of the fitted atomic model were deposited in the PDB under the accession codes 6Z10 (structure A) and 6Z1I (structure B). The cryo-EM data were deposited on EMPIAR with the accession code EMPIAR-10457. The datasets and materials used during the current study are available from the corresponding author on reasonable request. Source data are provided with this paper.

## Acknowledgements

We would like to thank the Deutsche Forschungsgemeinschaft for funding of the Research Unit FOR 2969, projects FA 456/27, HA 7138/3, HE 8472/1-1, HU 2400/1-1, SCHO 1364/2-1. We are grateful to Prof. Dr. Bernd Reif (Technical University Munich) for helpful discussions and to the students who participated in the practical course “Protein Biochemistry and Structural Biology” in the winter semester 2019, for assisting in the manual picking of the fibril segments. All cryo-EM data were collected at the European Molecular Biology Laboratory, Heidelberg (Germany), funded by iNEXT (Horizon 2020, European Union).

## Contributions

L.R. and J.B. carried out experiments. L.R, J.B., S.H., C.H., A.B., M.S. and M.F. analyzed data. U.H. and S.O.S. contributed tools and reagents. M.F. designed research. L.R. and M.F. wrote the paper.

