## Supplementary Information for "Cryo-EM reveals structural breaks in a patient-derived amyloid fibril from systemic AL amyloidosis"

L. Radamaker et al.

**Supplementary Figure 1**

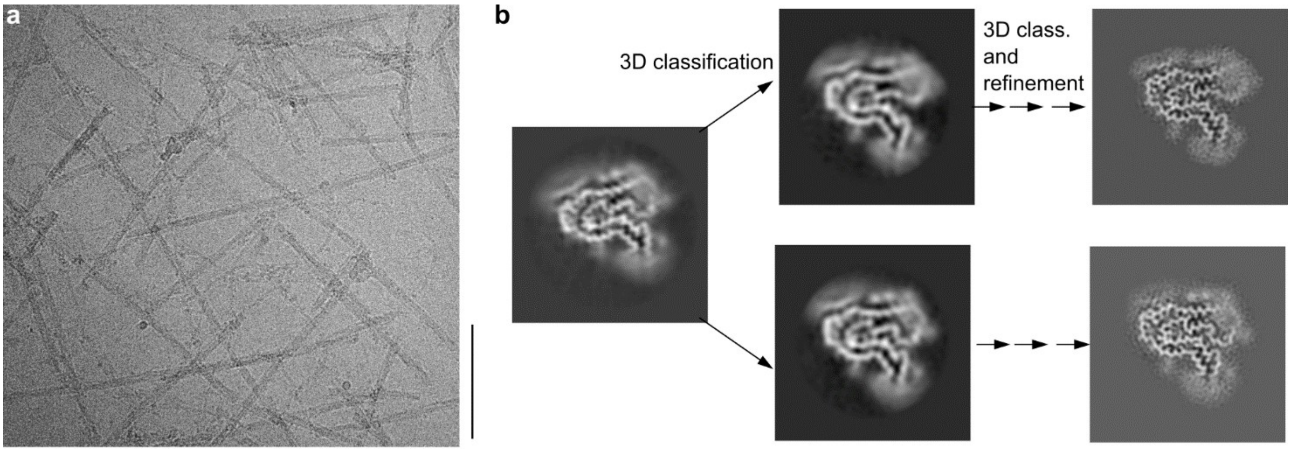

**Supplementary Fig. 1**

**Work flow of steps of the reconstruction process.**

(a) Cryo-EM image of the extracted fibrils. Scale bar: 100 nm. (b) Schematic illustration of the workflow. Based on an initial reconstruction, 3D classification separated two subsets exhibiting different density paths. Through several further classification and refinement steps, the final reconstructions were obtained.

**Supplementary Figure 2**

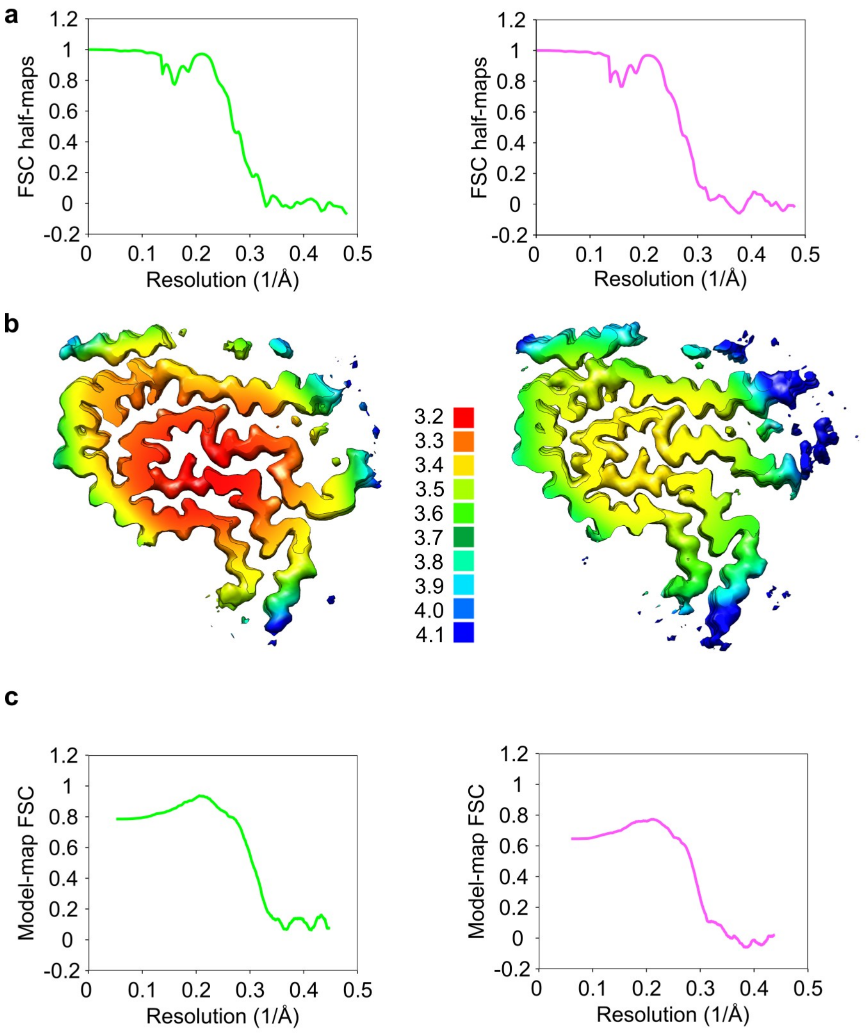

**Supplementary Fig. 2**

**Map resolution and model fit.**

(a) FSC curves for fibril structures A (green, left) and B (magenta, right). (b) Local resolution maps for A (left) and B (right). (c) Model-map FSC curves for models of conformation A (green, left) and B (magenta, right)

**Supplementary Figure 3**

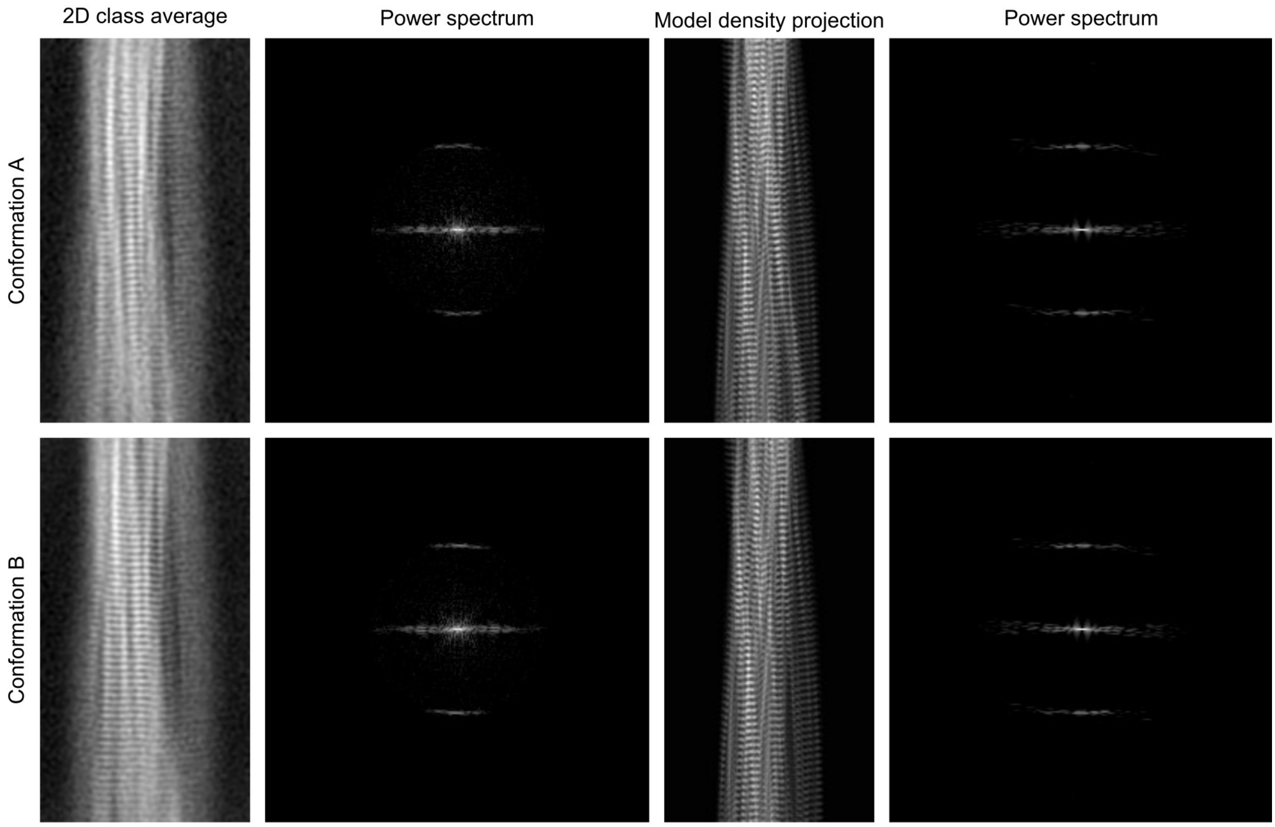

**Supplementary Fig. 3**

**Comparison of the molecular models with 2D class averages.**

For the two conformations A and B respectively, 2D class averages and model density projections corresponding to the same fibril regions are shown, with their respective power spectra. The length of the fibril section is approximately 266 Å.

**Supplementary Figure 4**

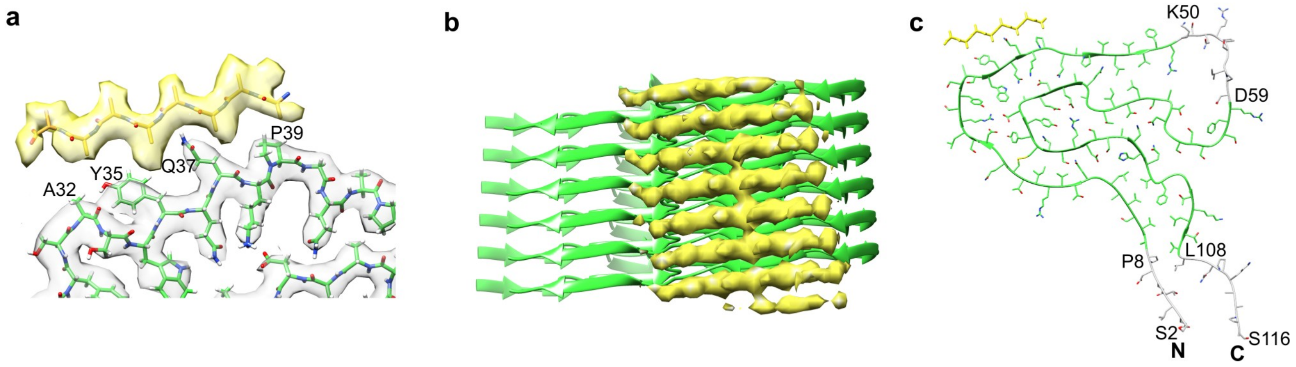

**Supplementary Fig. 4**

**Defined extra density decorating the fibril surface.**

(a) Enlarged view of the defined extra density (yellow) seen in structure A (green), overlayed with an 8-residue poly-L-alanine chain in parallel β-sheet conformation. (b) Side view of the defined extra density (yellow) next to the molecular model of a stack of fibril proteins in conformation A (ribbon diagram, green) showing the rise along the z-axis as well as the offset between the extra density and the fibril protein model. (c) Ribbon diagram including side chains of a single fibril protein layer in conformation A (green) showing all residues in the fibril protein including the disordered N- (Ser2-Pro8) and C-terminus (Leu108-Ser116) and the central disordered region (Lys50-Asp59) marked in gray, as well as the 8-residue poly-L-alanine chain as in (a) (yellow), representing the defined extra density. The gray segments are depicted in arbitrary conformation.

**Supplementary Figure 5**

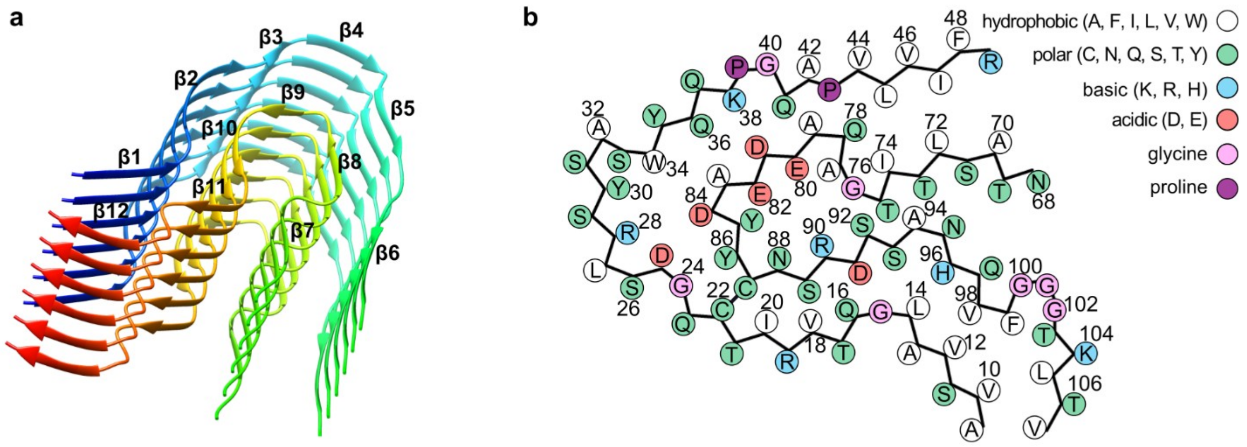

**Supplementary Fig. 5**

**Structural features of fibril protein conformation B.**

(a) Ribbon diagram of a stack of six fibril proteins in conformation B. (b) Schematic representation of the amino acid positions in conformation B. See Fig. 2b, c for the corresponding images showing conformation A.

**Supplementary Figure 6**

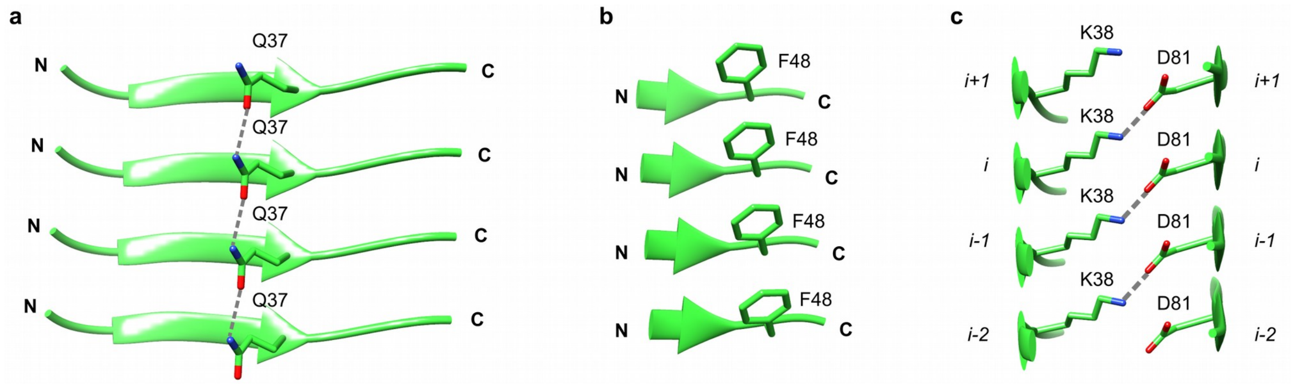

**Supplementary Fig. 6**

**Side chain-side chain interactions across the layers of the fibril model.**

(a) Stack of four fibril protein layers showing a polar ladder formed by Gln37 residues. (b) Four Phe48 residues in a configuration allowing for π-π (aromatic) stacking. (c) Electrostatic interactions between Lys38 of layer *i* and Asp81 of layer *i+1* demonstrating the interlocking of fibril protein layers. The images are based on a model of fibril structure A.

**Supplementary Figure 7**

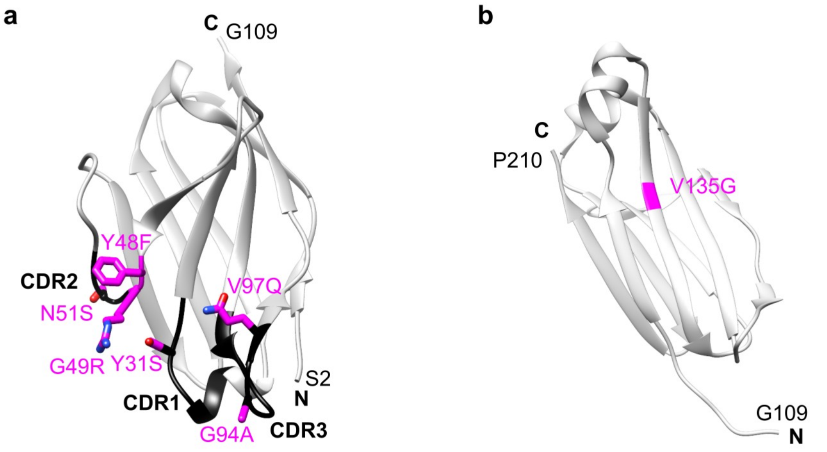

**Supplementary Fig. 7**

**Location of the FOR005 mutations within the natively folded LC domains.**

(a) Location of the mutations within the structure of the refolded FOR005 fibril protein (PDB: 5L6Q)[1]. (b) Location of the Val135Gly mutation, colored in magenta, in the $C_L$ domain of a homologous LC, containing an *IGLC2* GL segment (adapted from PDB: 4EOW[2,], see methods for details). CDRs are colored black, mutations magenta.

### Supplementary Figure 8

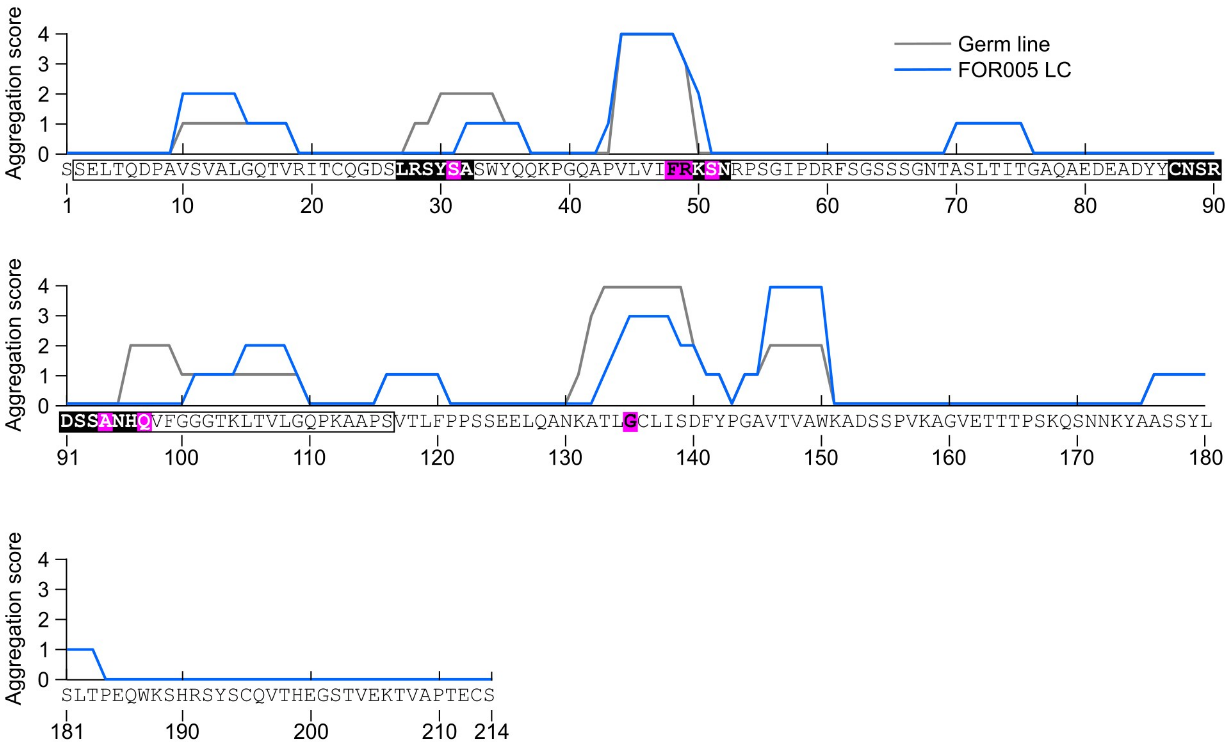

**Supplementary Fig. 8**

**Aggregation score of the FOR005 precursor LC and of the corresponding GL protein sequence.**

Aggregation score of the FOR005 LC precursor protein sequence (blue) and of the corresponding GL protein sequence (gray). The sequence on the x-axis is the FOR005 LC precursor protein sequence. The box denotes the range of the fibril protein (Ser2-Ser116). The black filled areas with white letters indicate the CDRs, residues deviating from the GL protein sequence are marked with magenta boxes. When located in a CDR, these residues are displayed in white: otherwise in black.

**Supplementary Figure 9**

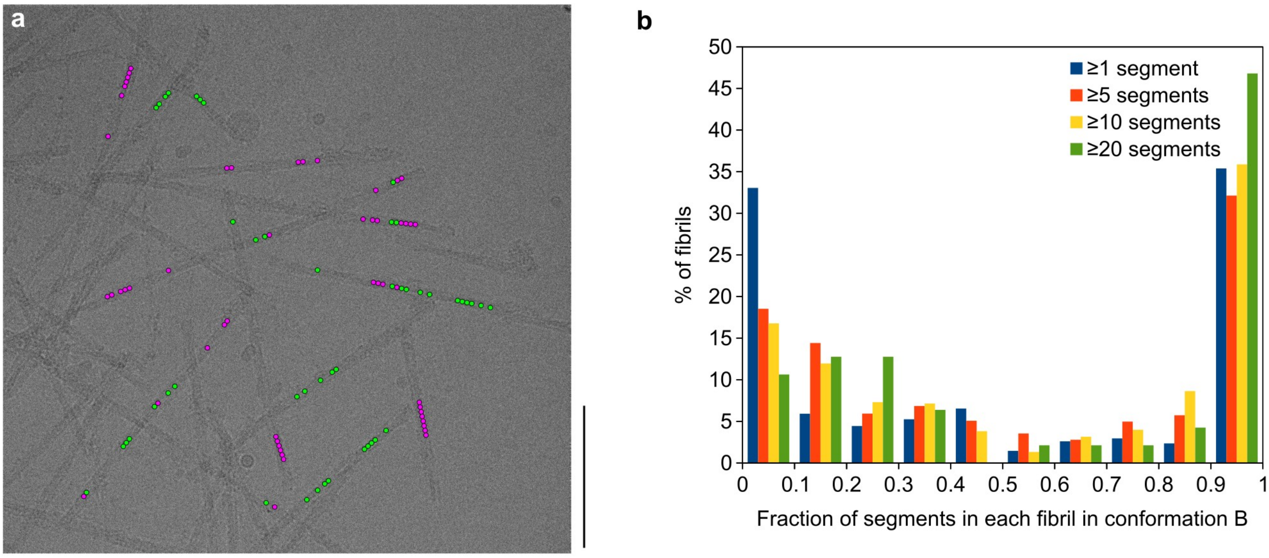

**Supplementary Fig. 9**

**Evidence for structural breaks**

(a) Representative cryo-EM micrograph showing the location of segments classified as fibril structure A (green) and fibril structure B (magenta). Scale bar: 100 nm. Figure 4a shows the same image, highlighting all the segments from the first round of 3D classification. (b) Histogram of the fraction of segments classified as fibril structure B, per fibril. Four different thresholds were chosen for the minimum total number of segments per fibril: a minimum of 1 segment (4,795 fibrils), 5 segments (1,970 fibrils), 10 segments (602 fibrils) and 20 segments (47 fibrils). The data set contained 11,003 segments for fibril structure A and 12,122 for fibril structure B. Figure 4b shows an analogous histogram including all the segments from the first round of 3D classification.

### Supplementary Figure 10

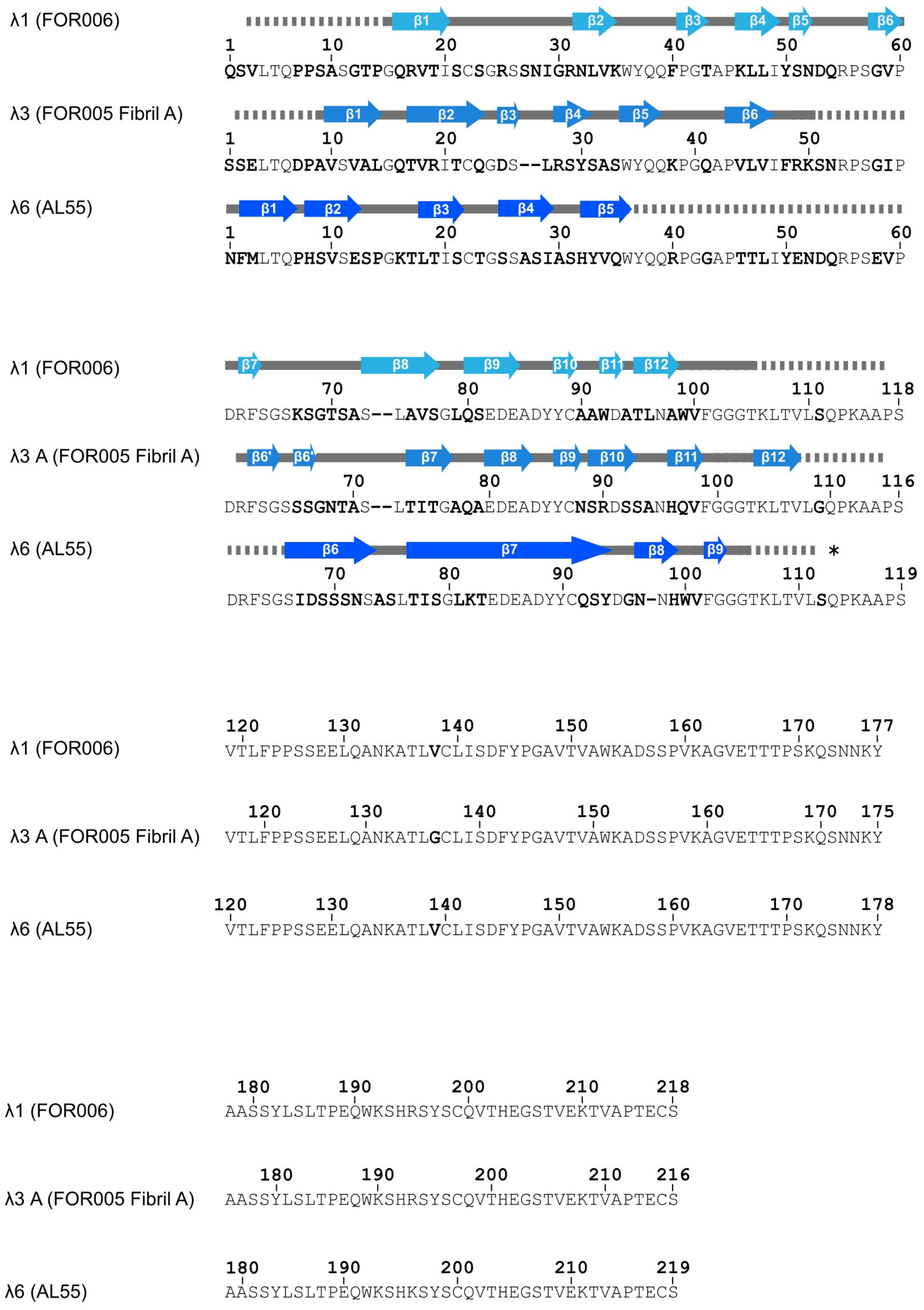

### Supplementary Fig. 10

**Sequence alignment and secondary structure comparison of ex vivo AL amyloid fibrils.**

Sequence alignment of ex vivo AL amyloid fibril LCs for which a cryo-EM structures is available, combined with a schematic representation of the secondary structure of these fibrils. Non-conserved residues are marked in bold. Arrows indicate β-strands. Continuous lines indicate ordered conformation, dotted lines indicate disordered segments. Star: no single truncation site reported. The definition of secondary structural elements follows the definition in the respective manuscripts (FOR006[3], FOR005, AL55[4]).

**Supplementary Table 1**

| | Fibril A (EMD-11031) (PDB 6Z10) | Fibril B (EMD-11030) (PDB 6Z1I) |
|---|---|---|
| **Data collection and processing** | | |
| Magnification | 130,000 | 130,000 |
| Voltage (kV) | 300 | 300 |
| Electron exposure (e– Å$^{-2}$) | 1 | 1 |
| Defocus range (μm) | 0.2 – 4.2 | 0.2 – 4.2 |
| Pixel size (Å) | 1.04 | 1.04 |
| Symmetry imposed | C1 | C1 |
| Initial particle images (no.) | 194,502 | 194,502 |
| Final particle images (no.) | 11,003 | 12,122 |
| Map resolution (Å) | 3.2 | 3.4 |
| FSC threshold (0.143) | | |
| Rise (Å) | 4.8 | 4.8 |
| Twist | -1.1 | -1.1 |
| Pitch (nm) | 155 | 155 |
| **Refinement** | | |
| Initial model used (PDB code) | De novo | PDB 6Z10 |
| Model resolution (Å) | | |
| FSC threshold 0.143 | 3.1 | 3.2 |
| FSC threshold 0.5 | 3.4 | 3.5 |
| Map CC (mask) | 0.76 | 0.73 |
| EMRinger ZScore | 5.09 | 5.97 |
| EMRinger Score | 2.88 | 3.59 |
| Model composition | | |
| Non-hydrogen atoms | 3,972 | 3,648 |
| Protein residues | 534 | 486 |
| Ligands | 0 | 0 |
| *B* factors (Å$^2$) | | |
| Protein | 73.24 | 72.43 |
| Ligand | - | - |
| R.m.s. deviations | | |
| Bond lengths (Å) | 0.009 | 0.009 |
| Bond angles (°) | 1.795 | 1.745 |
| Validation | | |

| | | |
|---|---|---|
| MolProbity score | 1.42 | 1.61 |
| Clashscore | 1.42 | 3.08 |
| Poor rotamers (%) | 0 | 0 |
| Ramachandran plot | | |
| Favored (%) | 91.76 | 91.34 |
| Allowed (%) | 8.24 | 8.66 |
| Disallowed (%) | 0 | 0 |
| B-factor after autosharpening (Å²) | 71.3 | 51.8 |
| Contour level | 6σ | 6σ |

**Supplementary Table 1.**

**Reconstruction and modeling statistics.**
